# Factorial Knockouts Distinguish Physical Necessity from Numerical Compensation in a PSI–LHCI Transport Model

**DOI:** 10.64898/2026.08.26.747189

**Authors:** Hai Zhang, Boyi Feng, Huan Tan, Yuqi Wang, Haoming Luo

## Abstract

Mechanistic interpretation of photosynthetic energy-transfer models requires more than reproduction of experimental observables: a model intended to support mechanistic claims should also respond consistently when its proposed functional organizations are removed. Here, we evaluate whether a calibrated PSI–LHCI transport surrogate encodes physically meaningful principles by applying a 2 × 2 factorial knockout framework that independently removes site-energy heterogeneity and coupling-strength heterogeneity in a 155-pigment network. All perturbations were evaluated using the same Full-model calibration without parameter refitting. Although the calibrated model reproduced a high excitation-trapping yield, eliminating either energetic or coupling heterogeneity unexpectedly improved its apparent transport performance. A strict zero-coupling control confirmed that coupling itself remained necessary for network-mediated reaction-center access, whereas the supplied organization of coupling strengths was not supported by the surrogate. An audit of the model inputs further identified peripheral localization of all lowest-energy states and effective coupling scales far above those used in structure-based chlorophyll Hamiltonians. These findings do not imply that native PSI favors flat energy landscapes or uniform couplings. Instead, they show that endpoint agreement alone does not validate a mechanistic interpretation of a pigment-network model. Factorial knockout analysis provides a falsification-oriented framework for separating physical necessity from proposed organization, diagnosing numerical compensation, and identifying the constraints required for more predictive models of PSI–LHCI energy transfer.

**Significance Statement:** Photosynthetic energy-transfer models are often judged by whether they reproduce a measured efficiency or lifetime. A successful fit, however, can hide incorrect internal physics when several parameters compensate for one another. We introduce a factorial knockout test that asks two separate questions: must pigment coupling exist, and does the modeled organization of pigment energies and coupling strengths actually improve transport? Applied to a calibrated PSI–LHCI surrogate, the test confirms that coupling is essential but shows that removing either modeled organization improves the fitted performance. Targeted controls trace this failure to peripheral low-energy assignments and an unrealistically large coupling scale. The framework turns an apparently successful simulation into a falsifiable mechanistic model and identifies which physical constraints must be repaired before biological design principles can be inferred.

## 1 Introduction

Photosynthetic energy conversion begins with absorption of solar radiation by pigment–protein complexes embedded in the thylakoid membrane. In oxygenic photosynthesis, photosystem II (PSII) oxidizes water and supplies electrons to the plastoquinone pool; electron transfer through cytochrome *b*_6_*f* and plastocyanin then delivers reducing equivalents to photosystem I (PSI), where a second photochemical excitation supports ferredoxin reduction and, ultimately, NADPH formation [1].

PSI is a particularly useful system for examining how a dense pigment network captures and transports excitation energy. In plants, the PSI core is surrounded by light-harvesting complex I (LHCI), and structurally resolved interfacial chlorophylls connect the peripheral antenna to the core [2, 3]. This pigment architecture can be represented as a structured excitation-transport network, in which individual chlorophylls act as energy-transfer sites connected by pigment–pigment electronic couplings. In the present PSI–LHCI transport surrogate, this architecture is represented as a network of 155 pigment sites connected by 409 pigment–pigment interactions, preserving the spatial organization required to examine how energetic and coupling heterogeneity influence excitation trapping. Excitation then migrates through this network before photochemical trapping at the P700 reaction center. Time-resolved measurements and kinetic analyses show rapid trapping on a tens-of-picoseconds scale and a high internal quantum efficiency, although the observed kinetics depend on antenna composition and excitation conditions. [4–8].

Two components of an effective excitonic Hamiltonian are central to such transport descriptions: pigment site energies and pigment–pigment electronic couplings. Site energies represent local excitation energies within heterogeneous protein environments, whereas couplings encode electronic communication determined by pigment separation, orientation, and transition-density structure [9–12]. Together with environmental broadening and relaxation, these quantities define the transfer-rate network used by reduced excitation-transport models.

The functional interpretation of the energetic landscape remains unresolved. Low-energy, farred chlorophyll states extend the absorption range of PSI–LHCI, but their population can slow photochemical trapping when escape requires thermal activation [13, 14]. Their effect is therefore location dependent. A low-energy state that remains well connected to the core can support efficient transfer, whereas a peripheral low-energy state can delay successful trapping [10, 13, 15]. Spectroscopic and structure-based studies further show that the lowest-energy LHCI states cannot always be represented as isolated scalar site energies: excitonic mixing and charge-transfer character contribute to the red-shifted states of Lhca complexes [16, 17].

Electronic coupling presents a related distinction. Nonzero coupling is required for excitation to migrate between pigment sites, but this necessity does not establish that the native allocation of stronger and weaker couplings is itself optimized for reaction-center trapping. Structure-based network models have generated valuable hypotheses about transport pathways, linker pigments, and excitation lifetimes [9–11]. Nevertheless, fitted endpoints may remain accurate when parameters compensate for an incomplete Hamiltonian, an inaccurate environmental model, or weakly identifiable parameter combinations [18, 19]. A mechanistic claim therefore requires more than agreement at a single calibrated endpoint [20, 21].

This concern motivates an intervention-based test. Previous kinetic-network studies have assessed photosynthetic design principles using perturbations, randomized networks, pathway statistics, and first-passage observables, notably in PSII supercomplex models [22, 23]. Such analyses ask whether a supplied organization performs differently from alternatives. The present study asks a complementary and more direct question: what happens when the proposed organization itself is removed while all calibrated parameters are held fixed?

We introduce a 2 × 2 factorial knockout that independently removes site-energy heterogeneity and coupling-strength heterogeneity from a calibrated 155-pigment PSI–LHCI surrogate. The four factorial conditions retain the same pigment sites, 409-edge topology, reaction-center definition, trapping and loss channels, and Full-model calibration. A separate *J* = 0 control tests whether coupling exists, whereas the factorial coupling knockout tests whether coupling strengths are organized. Terminal charge-separation yield, reaction-center access, and conditional first-passage time are evaluated without recalibration.

Unexpectedly, removing either energetic or coupling-strength heterogeneity improves the surrogate at its calibrated point. We interpret this outcome as a failed mechanistic intervention test rather than as evidence that native PSI prefers flat energies or uniform couplings. Targeted controls and parameter audits show that a local reaction-center energy offset is beneficial, whereas the supplied peripheral low-energy states and an extreme effective coupling scale generate compensating behavior. The factorial knockout thus reframes the model from an apparent description of native design into a diagnostic object whose failures identify missing physical structure.

## 2 Results

### 2.1 A factorial knockout separates existence from organization

The four factorial conditions independently manipulated site-energy heterogeneity and coupling-strength heterogeneity while retaining the same 155 sites and 409-edge topology (Figure 1). Full retained both supplied distributions. Energy-KO assigned a common site energy while retaining the Full coupling matrix. Coupling-KO retained the Full site energies but replaced the heterogeneous occupied-edge magnitudes with a common root-mean-square magnitude. Double-KO combined both interventions. No knockout was recalibrated.

**Figure 1.**
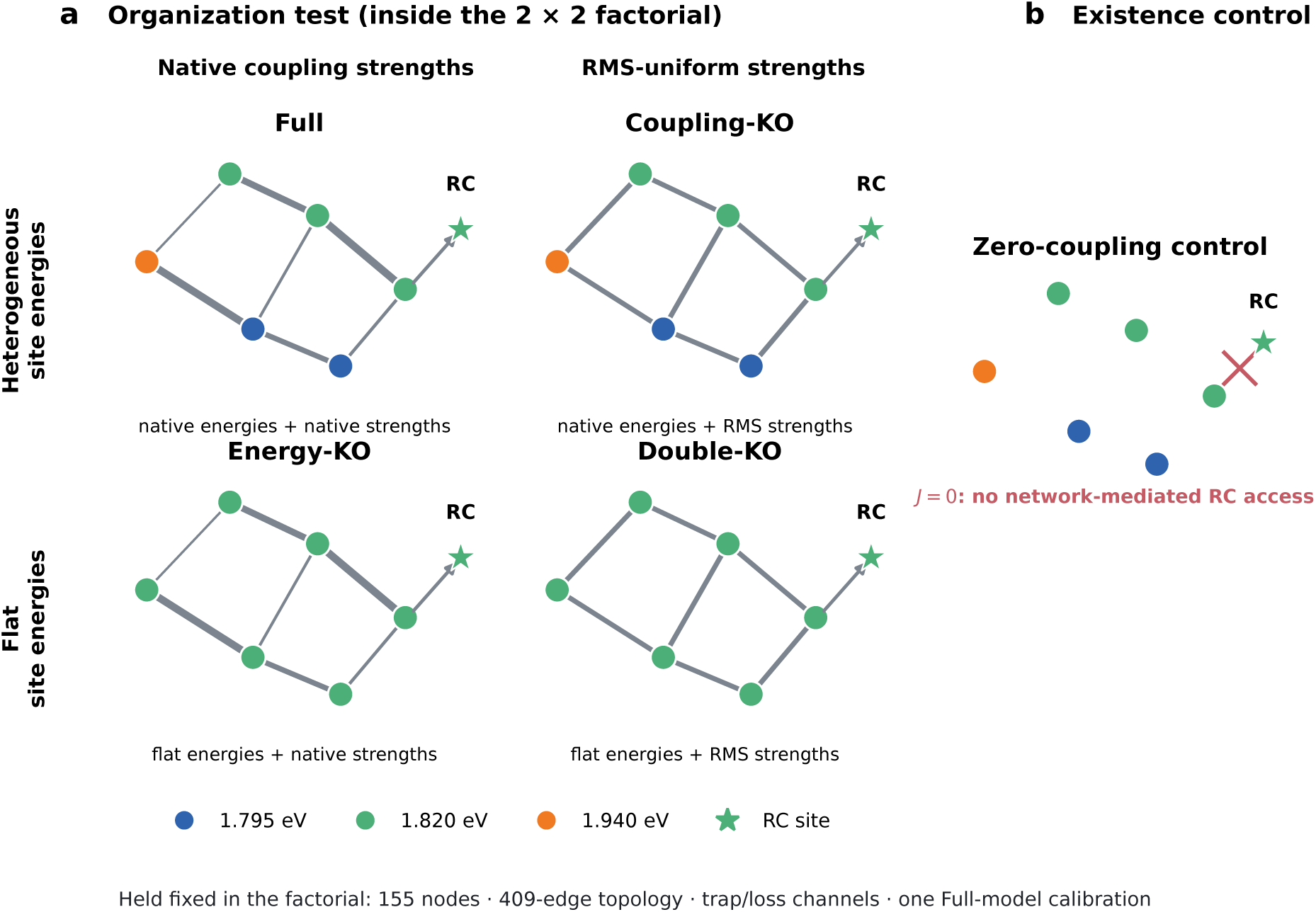
A factorial design separates existence from organization. **a**, The 2 × 2 organization knockout independently removes site-energy heterogeneity and coupling-strength heterogeneity. All conditions retain the same 155 pigment sites, 409-edge topology, trapping and loss channels, and Full-model calibration. Coupling-KO preserves total squared coupling magnitude. **b**, The strict zero-coupling control is external to the factorial because it asks whether transport exists, not whether coupling strengths are organized.

This design tests organization, not existence. Coupling-KO preserves a connected transfer network and therefore cannot establish whether coupling is necessary. In the separate *J* = 0 control, removal of every pigment–pigment coupling abolished network-mediated access to the reaction-center set. Coupling is therefore essential for transport in the surrogate. The relevant factorial question is narrower: does the supplied distribution of strong and weak couplings improve transport relative to a topology-preserving homogenization? Keeping these questions separate prevents a failure of coupling-strength organization from being misreported as dispensability of coupling itself.

### 2.2 The reference energetic landscape lacks a simple reaction-center-directed funnel

Before interpreting knockout performance, we audited the supplied site-energy map. The Full input contained three discrete energy classes. All 23 pigments in the lowest-energy class (1.795 eV) were assigned to peripheral LHCI, whereas both reaction-center sites occupied the intermediate class (1.820 eV; Figure 2). Thus, the supplied landscape was not a monotonic energetic funnel toward the reaction center.

**Figure 2.**
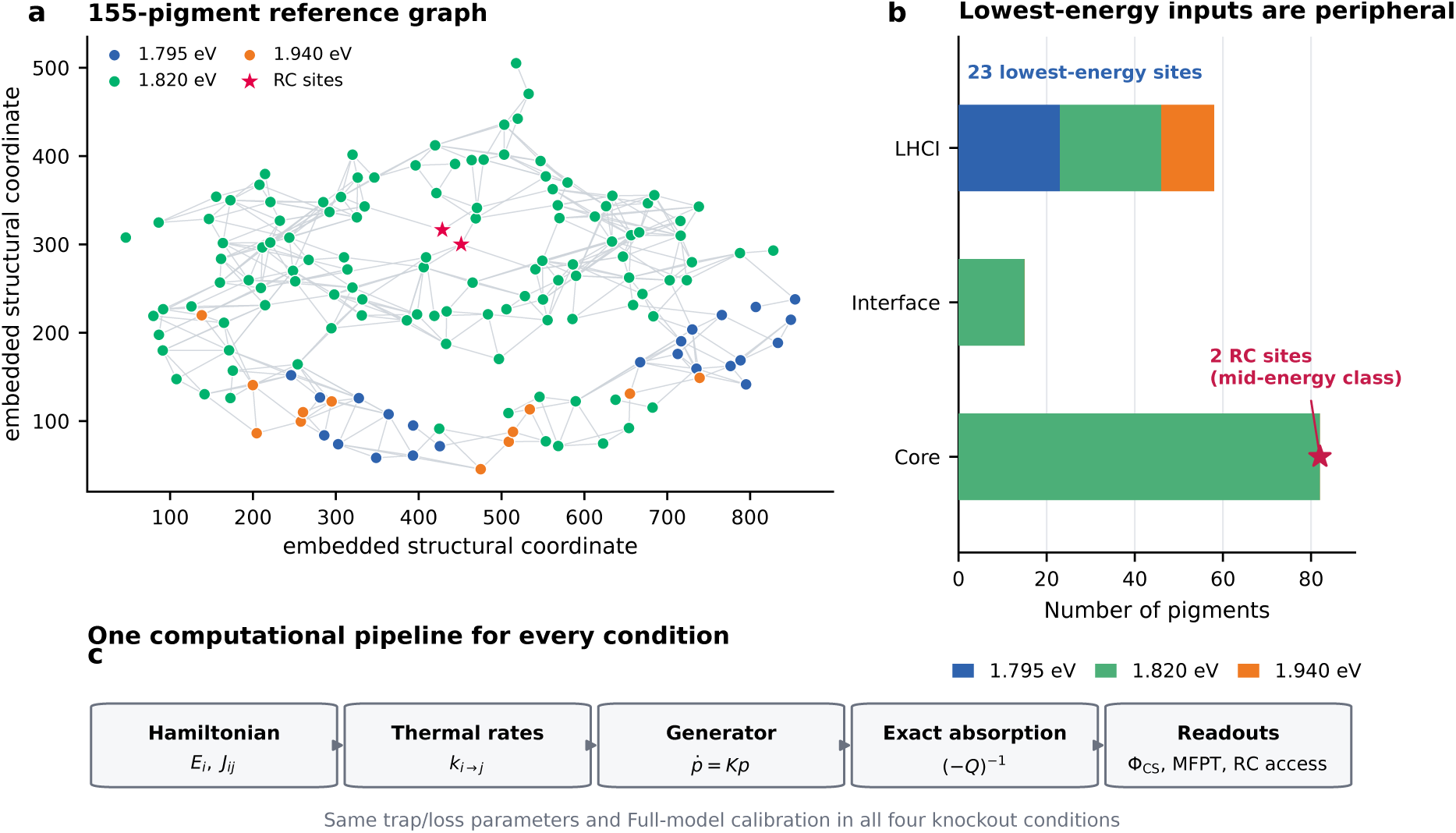
The reference energetic landscape lacks a simple reaction-center-directed funnel. **a**, Embedded representation of the 155-pigment reference graph, colored by the three supplied site-energy classes; stars denote the two reaction-center sites. **b**, Site-energy counts by coarse structural region. All 23 pigments in the lowest-energy class are assigned to peripheral LHCI, whereas both reaction-center sites occupy the intermediate class. **c**, The same Hamiltonian-to-rate-to-generator-to-absorption pipeline is applied to every knockout condition.

This observation does not establish that low-energy LHCI states are biologically misplaced. Real PSI–LHCI contains red-shifted states, and their transport consequences depend on microscopic origin, coupling, and structural context [10, 13–15]. The audit instead identifies a limitation of the present three-level surrogate: the scalar energy assignment places its deepest states exclusively at the periphery and does not represent the exciton–charge-transfer character associated with experimentally characterized red forms [16, 17]. The same Hamiltonian-to-rate-to-generator pipeline was applied to all conditions, so subsequent differences arise from interventions on this supplied representation rather than from different numerical treatments.

### 2.3 Removing energetic or coupling-strength heterogeneity improves the calibrated surrogate

At the calibrated point, the Full model reproduced the target charge-separation yield, F_CS_ = 0.965000, with reaction-center access *A*_RC_ = 0.988019, conditional CS-MFPT = 34.986 ps, and mean transient lifetime = 43.750 ps (Table 1; Figure 3). Despite this high endpoint performance, neither organization knockout impaired the surrogate.

**Table 1.** Primary knockout endpoints at the Full-model calibration point. Every knockout uses the Full calibration without parameter refitting.

| Condition | CS yield | CS-MFPT (ps) | RC access | Lifetime (ps) |
| --- | --- | --- | --- | --- |
| Full | 0.965000 | 34.986 | 0.988019 | 43.750 |
| Energy-KO | 0.973472 | 25.155 | 0.993460 | 33.160 |
| Coupling-KO | 0.970224 | 29.242 | 0.993458 | 37.221 |
| Double-KO | 0.973475 | 25.150 | 0.993464 | 33.156 |

**Figure 3.**
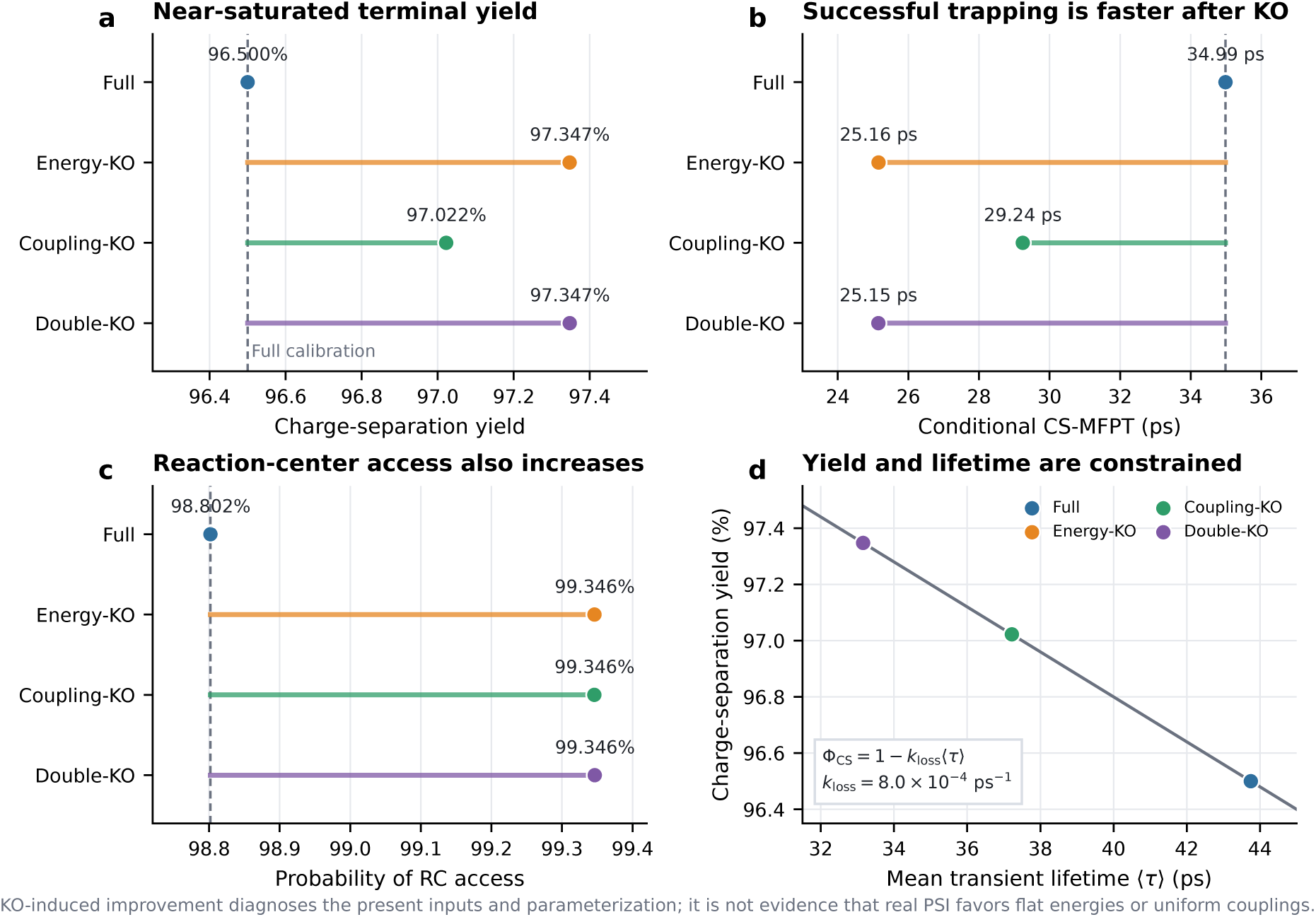
Organization knockouts improve the calibrated surrogate. **a**, Terminal charge-separation yield. **b**, Conditional charge-separation mean first-passage time (CS-MFPT). **c**, Probability of reaching a reaction-center site. Dashed lines in **a–c** mark the Full-model value. **d**, Terminal yield and mean transient excitation lifetime obey F_CS_ = 1 − *k*_loss_ ⟨*t*⟩ with *k*_loss_ = 8.0 × 10^−4^ ps^−1^ and therefore are not independent validations.

Energy-KO increased the yield to 0.973472, increased reaction-center access to 0.993460, and shortened conditional CS-MFPT to 25.155 ps. Coupling-KO likewise increased the yield to 0.970224, increased access to 0.993458, and shortened conditional CS-MFPT to 29.242 ps. Double-KO produced nearly the same outcome as Energy-KO: yield 0.973475, access 0.993464, and conditional CS-MFPT 25.150 ps.

These results distinguish coupling necessity from coupling-strength organization. The *J* = 0 control shows that transfer paths are required, whereas Coupling-KO shows that the native allocation of stronger and weaker edge weights is not supported as performance enhancing at the calibrated point. Similarly, the Energy-KO result is not evidence that biological PSI should possess a flat energy landscape. It shows that the energetic organization encoded by this particular input penalizes the model observables used in the calibration.

### 2.4 Targeted controls identify opposing energetic effects

To determine why Energy-KO succeeded, we decomposed the intervention into targeted energetic controls (Figure 4). Removing the coarse directionality of the supplied landscape increased yield from 0.9650 to 0.9721 and shortened conditional CS-MFPT from approximately 35.0 to 26.9 ps. Flattening the non-reaction-center antenna while retaining the reaction-center offset produced the highest yield (0.9773) and the shortest conditional CS-MFPT (20.4 ps).

**Figure 4.**
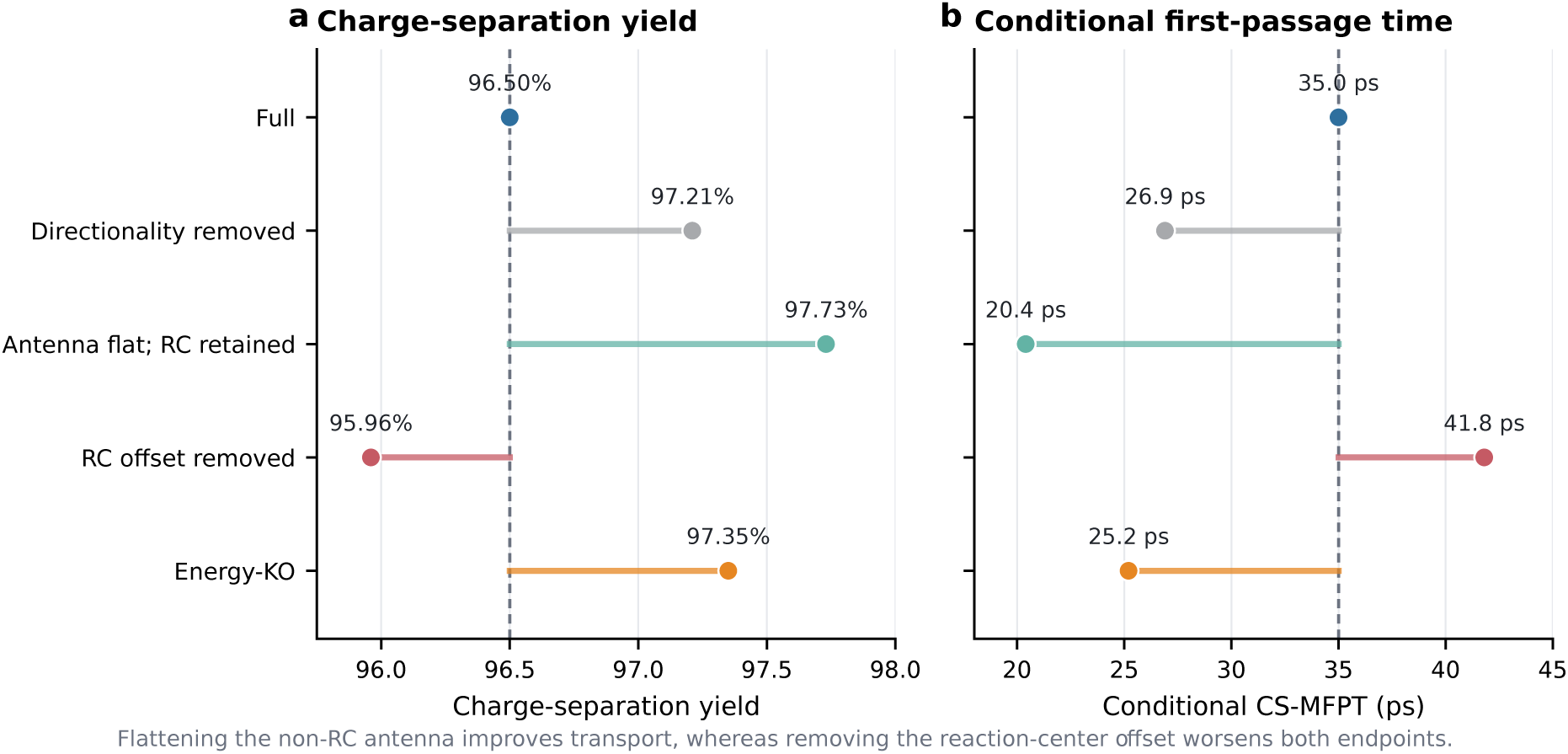
Targeted controls decompose the energy knockout. **a**, Charge-separation yield. **b**, Conditional CS-MFPT. Flattening the non-reaction-center antenna while retaining the reaction-center offset improves both outcomes, whereas removing only the reaction-center offset lowers yield and slows successful trapping.

The reaction-center offset itself was beneficial. Removing only that offset lowered the yield to 0.9596 and lengthened conditional CS-MFPT to 41.8 ps. Complete Energy-KO gave an intermediate response (yield 0.9735; conditional CS-MFPT 25.2 ps) because it removed both the detrimental peripheral organization and the beneficial local reaction-center bias.

The relevant inference is therefore not that energetic organization is generically harmful. Rather, this coarse input combines two opposing effects: a useful local reaction-center bias and peripheral low-energy assignments that impede trapping. This location dependence is consistent with experimental and theoretical work showing that red states can either delay trapping through thermally activated escape or remain efficient when their structural connections support transfer toward the core [10, 13–15].

### 2.5 Endpoint yield and transport speed need not rank models identically

For a constant loss rate in this absorbing model, terminal charge-separation yield and mean transient lifetime obey

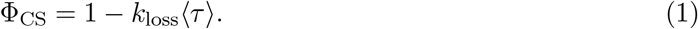

With *k*_loss_ = 8.0 × 10^−4^ ps^−1^, fitting Φ_CS_ = 0.965 fixes ⟨*τ*⟩ = 43.75 ps. These quantities are algebraically linked rather than independent validations. In addition, near-saturated yield compresses kinetic differences: substantial changes in first-passage time can produce only small changes in terminal yield. Treating both quantities as independent calibration successes would overstate the evidence supplied by the endpoint fit, a general concern in sloppy or discrepancy-containing mechanistic models [18, 19].

The coupling-scale scan made this distinction explicit (Figure 5). At *J*_max_ = 120 cm^−1^, Energy-KO increased terminal yield relative to Full but lengthened conditional CS-MFPT from approximately 207 to 268 ps. Thus, a condition can win according to eventual trapping probability while losing according to the speed of successful trapping. Near the calibrated coupling scale, the yields of all four models approached saturation, whereas kinetic differences remained visible. Endpoint yield alone therefore cannot rank the transport organizations.

**Figure 5.**
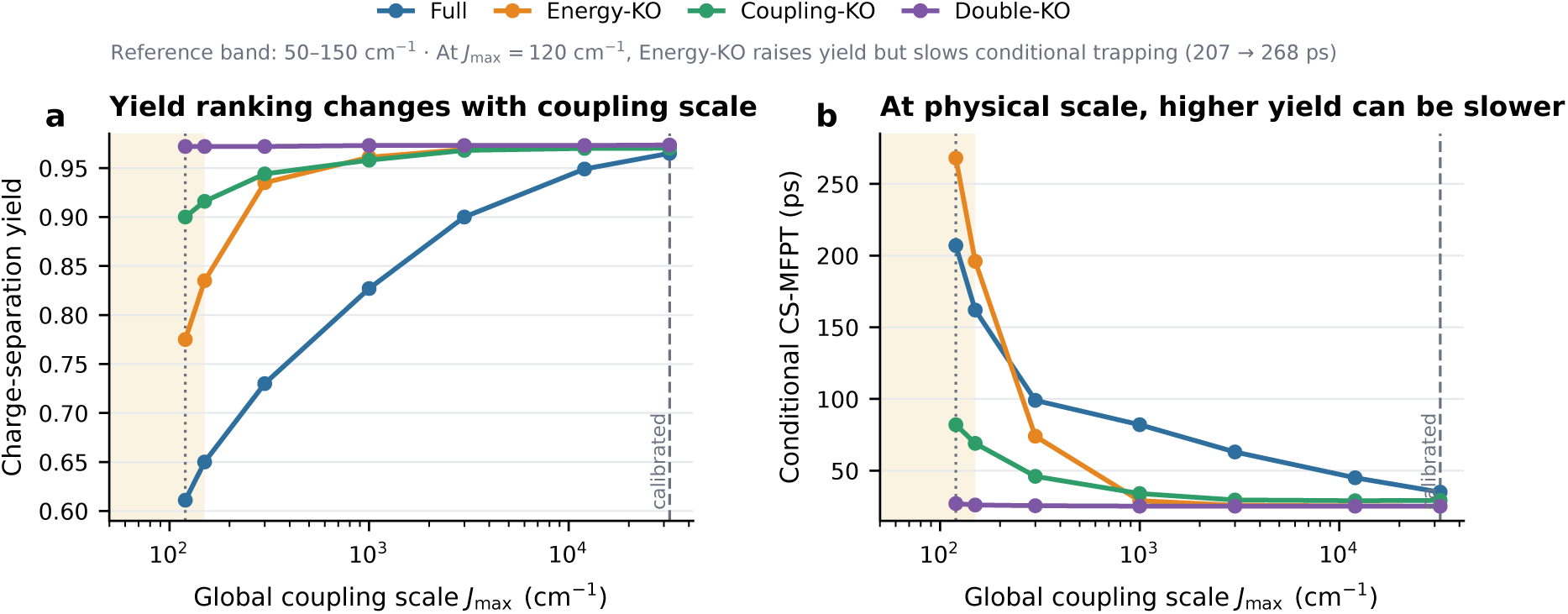
Coupling-scale sensitivity exposes a yield–speed tradeoff. **a**, Charge-separation yield across the global coupling scale *J*_max_. **b**, Conditional CS-MFPT over the same scan. The shaded band indicates a representative 50–150 cm^−1^ chlorophyll-coupling range, the dotted line marks *J*_max_ = 120 cm^−1^, and the dashed line marks the calibrated scale. At *J*_max_ = 120 cm^−1^, Energy-KO increases terminal yield but slows successful trapping.

### 2.6 The calibrated coupling scale prevents a native-design interpretation

An internal scale audit showed that the effective coupling magnitudes used at calibration were not compatible with a direct structural interpretation (Figure 6). The reference comparison used *J*_max_ = 120 cm^−1^, within scales commonly encountered in structure-based chlorophyll Hamiltonians [9–12]. By contrast, Coupling-KO assigned every occupied edge a magnitude of 5,122 cm^−1^, and the calibrated maximum scale was approximately 3.2 × 10^4^ cm^−1^. The 50–150 cm^−1^ band shown in Figures 5 and 6 is a representative comparison range rather than a universal hard upper bound; nevertheless, the knockout and calibrated values exceed it by orders of magnitude.

**Figure 6.**
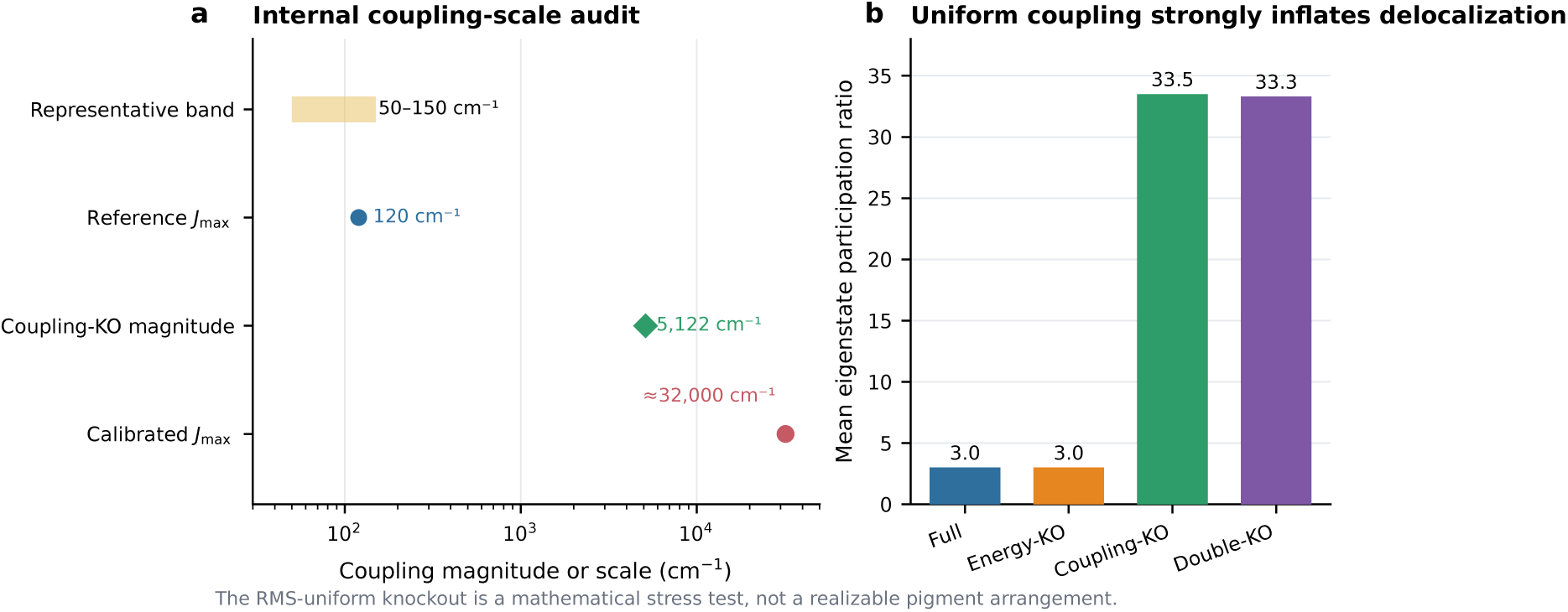
The calibrated coupling scale precludes a native-design interpretation. **a**, Internal scale audit comparing a representative 50–150 cm^−1^ band, the reference *J*_max_ = 120 cm^−1^, the 5,122 cm^−1^ RMS-uniform Coupling-KO magnitude, and the approximately 3.2 × 10^4^ cm^−1^ calibrated maximum scale. **b**, Mean Hamiltonian eigenstate participation ratio. Uniform coupling strongly increases apparent delocalization in Coupling-KO and Double-KO.

Uniform coupling also inflated the mean Hamiltonian eigenstate participation ratio from approximately 3.0 in Full and Energy-KO to 33.5 and 33.3 in Coupling-KO and Double-KO, respectively. The resulting apparent delocalization is a consequence of the mathematical intervention at an extreme scale and should not be interpreted as a realizable alternative pigment arrangement. The fitted coupling scale instead behaves as a compensatory parameter that helps the surrogate reproduce the endpoint despite deficiencies elsewhere in the physical description.

## 3 Discussion

### 3.1 Knockout improvement is a model diagnosis, not a biological optimum

The central result is not that real PSI should remove energetic structure or homogenize its couplings. It is that the calibrated surrogate does not respond to these interventions in the way required for a strong native-design interpretation. A model that assigns functional importance to a particular organization should normally lose the associated function when that organization is removed, provided that the intervention preserves the remaining quantities needed for a fair comparison. Here, both organization knockouts improve the calibrated observables. The intervention therefore falsifies the claim that the supplied heterogeneities are responsible for the fitted transport performance.

This distinction is important because endpoint reproduction and mechanistic validity answer different questions. Calibration asks whether some parameter combination can reproduce selected observations. Intervention asks whether the fitted internal structure carries the proposed causal role. Sloppy sensitivities and model discrepancy make it possible for an inaccurate or incomplete representation to fit an endpoint through compensating parameters [18,19]. A knockout that improves performance is therefore informative: it exposes where the fitted model relies on compensation rather than supporting the original mechanistic interpretation [20, 21].

### 3.2 Existence and organization must be tested separately

The zero-coupling and Coupling-KO controls answer different questions. Setting *J* = 0 eliminates excitation migration and confirms that coupling existence is necessary for reaction-center access. Coupling-KO preserves paths and total squared coupling magnitude but erases the allocation of stronger and weaker edges. Its improved performance therefore does not make coupling dispensable; it shows only that the supplied pattern of coupling strengths is not demonstrated to be advantageous within this surrogate.

This separation generalizes beyond PSI. Ablation studies often conflate deletion of a mechanism with deletion of its heterogeneity or architecture. A factorial construction makes the preserved and removed quantities explicit and allows interaction terms to reveal whether energetic and coupling organizations depend on one another. The approach complements randomization and network-entropy analyses developed for photosynthetic transport [22, 23] while placing stronger emphasis on falsification of a fixed calibrated model.

### 3.3 Energetic organization is local and multidimensional

The targeted controls show that a single label such as “energy funnel” is too coarse. Retaining the reaction-center offset while flattening the non-reaction-center antenna gives the best outcome, whereas removing only the reaction-center offset is detrimental. Energetic organization therefore contains at least two functionally opposing components in the present input. The beneficial local bias is hidden by peripheral low-energy states that delay trapping.

The scalar-site representation also omits physics known to matter for LHCI red forms. Experimental and theoretical studies attribute these states to excitonic interactions, protein electrostatics, and charge-transfer mixing rather than to an isolated low-energy pigment alone [16, 17]. A predictive revision should consequently replace the three-level assignment with a structurally localized Hamiltonian that represents the relevant excitonic and charge-transfer manifolds and tests whether low-energy states are positioned and coupled consistently with measured spectra and trapping kinetics [10, 15].

### 3.4 Near-saturated yield conceals kinetic and scale failures

The Full yield of 0.965 appears impressive in isolation, but it is a weak discriminator near saturation. In this model, yield and mean lifetime are linked by Eq. 1; fitting one fixes the other when the loss rate is held constant. The coupling-scale scan further shows that yield and conditional first-passage time can rank conditions differently. A robust calibration should therefore include observables that constrain distinct aspects of the dynamics, such as wavelength-resolved lifetimes, antenna-to-core transfer components, reaction-center arrival distributions, and spectroscopic line shapes [5–8].

The scale audit is even more restrictive. Coupling-KO and the calibrated Full model operate at effective magnitudes orders of magnitude above the representative structure-based range. At such scales, the large participation ratio and rapid transport are mathematical properties of the rescaled Hamiltonian, not evidence for a realizable chlorophyll arrangement. The global coupling scale should therefore be constrained before calibration, rather than allowed to compensate freely for deficiencies in site energies, environmental relaxation, or trapping kinetics.

### 3.5 Physical constraints for a predictive next-generation model

The intervention results suggest four concrete requirements. First, site energies should be derived from or constrained by structure-specific spectroscopy rather than assigned only as three coarse classes. Second, the lowest-energy LHCI states should include exciton–charge-transfer character where supported by experiment. Third, electronic couplings should be calculated from pigment geometry and transition densities and restricted to a physically defensible range [12]. Fourth, calibration should use multiple independent kinetic and spectroscopic observables, with out-of-sample interventions or perturbations reserved for validation.

These changes would alter the interpretation of factorial knockouts. A physically constrained model might show that some energetic or coupling organization is genuinely beneficial, that only local motifs matter, or that performance is robust over a broad ensemble rather than uniquely optimized. Any of these outcomes would be more informative than a high fitted endpoint alone because the conclusion would survive explicit attempts to remove the proposed mechanism.

### 3.6 Limitations

The conclusions apply to one calibrated 155-pigment surrogate and not directly to native PSI–LHCI. The supplied energy map uses three scalar classes and does not resolve microscopic red-state composition. Coupling-KO is deliberately a mathematical stress test; at the calibrated scale it is not a plausible pigment arrangement. The representative 50–150 cm^−1^ comparison band is not a universal upper bound, and physical couplings depend on pigment pair, convention, and electronic-structure method. Finally, the analysis evaluates a fixed rate-based transport model and therefore does not quantify uncertainty associated with alternative environmental spectral densities or non-Markovian dynamics. These limitations strengthen, rather than weaken, the main inference: the present results diagnose missing physical constraints in the surrogate and should not be promoted into biological optimality claims.

## 4 Methods

### 4.1 Pigment-network representation

The working PSI–LHCI surrogate analyzed here contained *N* = 155 chlorophyll sites assigned to LHCI, the antenna–core interface, the PSI core, and the reaction-center region. Chlorophyll sites were represented as nodes, and each unique unordered pigment pair (*i, j*), with *i* < *j*, was treated as an occupied edge when its modeled electronic coupling satisfied *J*_*ij*_ */=* 0. Under this definition, the working network contained *M* = 409 occupied edges. The regional architecture was motivated by high-resolution structures of plant PSI–LHCI [2, 3], whereas the pigment-network abstraction follows established structure-based treatments of photosynthetic excitation transport [9, 11].

Each site *i* was assigned an energy *E*_*i*_, and each occupied pair (*i, j*) was assigned a coupling *J*_*ij*_. The one-excitation Hamiltonian was

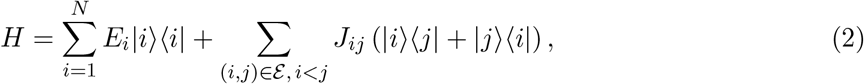

where ℰ is the occupied-edge set. Site energies were expressed in electronvolts in the supplied input, and effective coupling magnitudes were reported in cm^−1^.

### 4.2 Transport generator and Full-model calibration

For specified *E* and *J*, the reference pipeline converted the Hamiltonian into thermally weighted interstate rates and assembled a continuous-time population generator *K*(*E, J*; ***θ***). The reduced treatment belongs to the Haken–Strobl–Reineker class of stochastic exciton-transport descriptions [24, 25]. Population dynamics obeyed

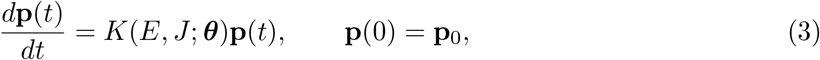

with irreversible reaction-center trapping and a state-independent loss channel. The same initial distribution, thermal-rate mapping, trap definition, loss rate, and numerical settings were used for every intervention.

The parameter vector ***θ***^∗^ was calibrated only for the Full model to reproduce Φ_CS_ = 0.965. The uniform loss rate was *k*_loss_ = 8.0 × 10^−4^ ps^−1^. All knockout and sensitivity calculations reused ***θ***^∗^ without refitting. This fixed-calibration design prevents altered models from recovering the target through a second round of parameter compensation.

### 4.3 Site-energy and coupling-strength knockouts

Energy-KO removed site-energy heterogeneity by assigning every site the Full-network arithmetic mean,

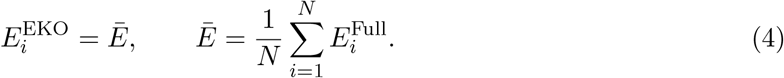

This preserved the number, positions, regional labels, and couplings of all pigments while eliminating energetic gradients and disorder.

Coupling-KO preserved the occupied-edge set but replaced each coupling magnitude with the Full-network root-mean-square value,

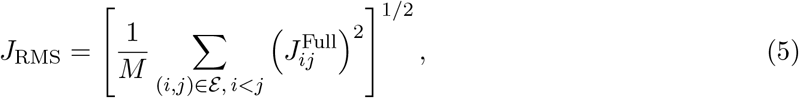

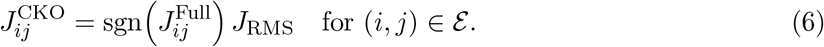

The transformation preserved topology, edge count, coupling-sign pattern, and total squared coupling magnitude while removing the distribution of strong and weak magnitudes. Double-KO combined the Energy-KO and Coupling-KO transformations.

### 4.4 Factorial contrasts and zero-coupling control

Let *X*_*ec*_ be an observable when energy heterogeneity *e* ∈ {0, 1} and coupling-strength heterogeneity *c* ∈ {0, 1} are absent or present. Full, Energy-KO, Coupling-KO, and Double-KO correspond to *X*_11_, *X*_01_, *X*_10_, and *X*_00_, respectively. Average main effects and the interaction were

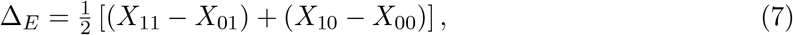

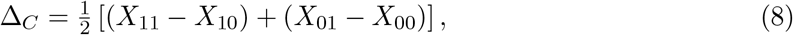

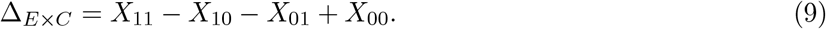

For yield and access, positive retention contrasts favor the retained organization. Because shorter first-passage time is favorable, time contrasts were interpreted with the opposite sign.

Coupling existence was tested outside the factorial by setting *J*_*ij*_ = 0 for all *i* ≠ *j*. The initial distribution for this control excluded reaction-center population so that measured access required transport through the pigment network.

### 4.5 Absorption probabilities and first-passage observables

The transient-state generator was augmented with absorbing charge-separation and loss states. If *Q* denotes the transient subgenerator, the fundamental matrix is

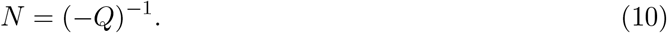

For initial row vector ***α***, the mean transient lifetime and terminal charge-separation yield were

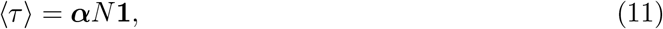

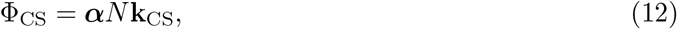

where **k**_CS_ contains absorption rates into the charge-separated state. With uniform loss and exhaustive charge-separation/loss outcomes, these expressions give Eq. 1.

Reaction-center access was evaluated by making the reaction-center set *R* absorbing before charge separation. If *f*_*R*_ (*t*) is first-arrival flux, access probability and conditional mean first-passage time were

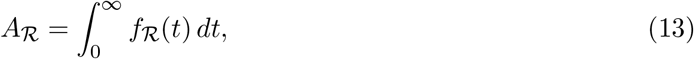

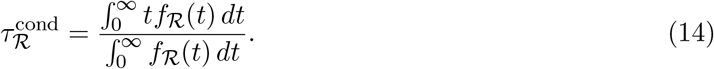

Conditional time was interpreted jointly with access probability because a short time among successful trajectories does not imply high overall accessibility.

### 4.6 Energetic and coupling-scale audits

Full-model site energies were mapped to the fixed pigment graph and ranked. Regional counts were computed for LHCI, antenna–core interface, core, and reaction-center sites. Targeted controls separately removed coarse directional assignment, flattened the non-reaction-center antenna while retaining the reaction-center offset, or removed only the reaction-center offset. Each control retained the Full coupling matrix and calibration.

Coupling-scale sensitivity was evaluated by varying the global coupling scale while holding all other parameters fixed. No scan point was recalibrated. Hamiltonian eigenstate delocalization was summarized by the participation ratio

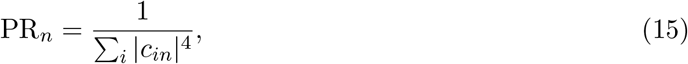

for normalized eigenstate |*ψ*_*n*_ ⟩ = ∑ _*i*_ *c*_*in*_ | *i* ⟩. The mean participation ratio was then compared among the four factorial conditions.

### 4.7 Statistical interpretation

The reported calculations are deterministic interventions on one calibrated surrogate. Results were therefore evaluated through effect sizes, factorial contrasts, and parameter sensitivity rather than replicate-based null-hypothesis tests. Numerical values were calculated from unrounded outputs.

## Data and Code Availability

The numerical inputs underlying the tables and figures, the four perturbed Hamiltonians, pigment-region annotations, and analysis code are available from the authors upon reasonable request. Public repository deposition is planned for a future release

## Acknowledgements

This work was developed during the 2026 PEBBLE BioFusion Workshop, held at Westlake University in Hangzhou, China, from 24 July to 4 August 2026.

